# De novo identification of toxicants that cause irreparable damage to parasitic nematode intestinal cells

**DOI:** 10.1101/852525

**Authors:** Douglas P. Jasmer, Bruce A. Rosa, Rahul Tyagi, Christina A. Bulman, Brenda Beerntsen, Joseph F. Urban, Judy Sakanari, Makedonka Mitreva

## Abstract

Efforts to identify new drugs for therapeutic and preventive treatments against parasitic nematodes have gained increasing interest with expanding pathogen omics databases and drug databases from which new anthelmintic compounds might be identified. Here, a novel approach focused on integrating a pan-Nematoda multi-omics data targeted to a specific nematode organ system (the intestinal tract) with evidence-based filtering and chemogenomic screening was undertaken. Based on *de novo* computational target prioritization of the 3,564 conserved intestine genes in *A. suum*, exocytosis was identified as a high priority pathway, and predicted inhibitors of exocytosis were tested using the large roundworm (*Ascaris suum* larval stages), a filarial worm (*Brugia pahangi* adult and L3), a whipworm (*Trichuris muris* adult), and the non-parasitic nematode *Caenorhabditis elegans*. 10 of 13 inhibitors were found to cause rapid immotility in *A. suum* L3 larvae, and five inhibitors were effective against the three phylogenetically diverse parasitic nematode species, indicating potential for a broad spectrum anthelmintics. Several distinct pathologic phenotypes were resolved related to molting, motility, or intestinal cell and tissue damage using conventional and novel histologic methods. Pathologic profiles characteristic for each inhibitor will guide future research to uncover mechanisms of the anthelmintic effects and improve on drug designs. This progress firmly validates the focus on intestinal cell biology as a useful resource to develop novel anthelmintic strategies.

**Author summary:** The intestinal cells of parasitic nematodes are not known to regenerate, therefore disruption of essential processes that cause irreparable damage to intestinal cells is expected to promote worm expulsion. To facilitate improved methods of therapy we need to better understand the basic intestinal cell and tissue functions of this critical organ. To that end have undertaken a comprehensive analysis of multi-omics omics data and identify and prioritize intestinal genes/pathways with essential functions and associated drugs and established a foundational model of the STH intestinal system using the large roundworm *Ascaris suum* to test and validate inhibitors of these functions. We found 10 inhibitors to impacted motility, and seven of those showed severe pathology and an apparent irreparable damage to intestinal cells. Furthermore, five inhibitors were effective against the three phylogenetically diverse parasitic nematode species, indicating potential for a broad spectrum anthelmintics. Our results firmly validate the focus on intestinal cell biology as a useful resource to develop novel anthelmintic strategies.

## Introduction

Nematode Infections in humans produce substantial mortality and morbidity, especially in tropical regions of Africa, Asia, and the Americas, leading to a number of important neglected tropical diseases. These pathogens include, but are not limited to, the intestinal worms referred to as soil transmitted helminths (STHs; mainly hookworms, ascarids, and whipworms) and filarial nematodes. STHs have high health impacts on the adult population as well as children by impairing growth and cognitive development, and causing anemia. STHs (with >0.5-1 billion infections for each of the three species listed) alone cause more morbidity than all parasitic diseases except malaria [1]. The filarial nematodes that cause, lymphatic filariasis and river blindness together affect hundreds of millions of people worldwide, particularly people living in impoverished conditions [2]. In addition, immune modulation by parasitic nematodes appears to interfere with immunity and vaccination against other pathogens [3–5]. Parasitic nematodes that infect livestock also reduce production of meat, milk and fiber; resources that play critical roles in the health and well-being of people, particularly in those regions most affected by nematode pathogens that directly impact human health.

As there are no vaccines, we must rely on behavior (hygienic practices), use of anthelmintics, and management of vector populations to limit health impacts of these pathogens. However, rapid re-infection after treatment which leads to temporary relief, the differential effectiveness of available anthelmintics and increasing emergence of nematode resistance against them [6] necessitate development of new therapeutics with possibly novel modes of action, broader parasite specificity and less susceptibility to the development of resistance.

The intestinal tract of parasitic nematodes is a high-priority drug target because 1) cells that comprise this organ system form a single polarized cell layer in direct contact with the outside host environment, 2) it performs essential functions associated with nutrient digestion and acquisition, reproduction, protection against environmental toxins [7, 8], and produces components implicated in interacting with the host immune system [9–11], and 3) intestinal cells are not known to regenerate, therefore disruption of essential processes that cause irreparable damage to intestinal cells is expected to promote worm expulsion. Many functions located at this host interface present targets for novel therapies to treat and control nematode infections, such as glycoproteins located on the apical intestinal membrane (AIM) that are effective targets for vaccines against human/animal parasitic nematodes [12, 13]. AIM proteins also stimulate host mucosal immune responses during infections [10] that may contribute to local immunity. Intestinal cells of some nematodes are also hypersensitive to benzimidazole (BNZ) anthelmintics [14, 15], which cause disintegration of intestinal cells, and the AIM is also a primary target for pore-forming crystal toxins produced by *Bacillus* spp. [16], which are effective against human/animal parasitic nematodes [17]. In each case, characteristics unique to intestinal cells appear to account for the anthelmintic effects related to these immunization, drug, and protein-toxin treatments.

To better understand the basic intestinal cell and tissue functions of this critical organ in parasitic nematodes and facilitate improved methods of therapy, we have established a foundational model of the STH intestinal system using *Ascaris suum*. With this model, we have identified 1) genes, proteins and predicted functions characterizing 10 different adult *A. suum* tissues (including the intestine) using microarrays [18] and RNA-seq [19], 2) transcripts, proteins and functions that are preferentially or constitutively expressed among three contiguous regions of the intestine [20], and 3) intestinal proteins that differentially localize to several intestinal cellular compartments by LC-MS/MS [21, 22]. This information was integrated with pan-Nematoda intestinal protein family databases [23] developed for the purpose of broad application of intestinal research to many different nematode pathogens. One intended use of these resources has been to predict intestinal cell targets and identify corresponding inhibitors for advancing anthelmintic research, the first test of which is reported here. We intersect results of interest from these multi-omics databases and demonstrate that using advanced computational, functional genomic and experimental screens, we can enable systematic and comprehensive identification of therapeutic targets and associated small molecule inhibitors on STHs (*A. suum* and *Trichuris muris*), the filarial nematode *Brugia pahangi*, and the non-parasitic nematode *C. elegans*. This long-term effort culminated in the identification of inhibitors and successful demonstration of activity against nematodes from across the phylum. Pathologic effects induced in intestinal cells and tissue illustrate an array of detrimental effects, including apparent irreparable damage, that is caused by several repurposed drugs and other inhibitors. Thus, this omics-based approach and focus on intestinal cells and tissue of parasitic nematodes provides a unique and powerful approach with application to identify new anthelmintic candidates, while also opening multiple avenues of future research on understanding global cellular responses of pathogens to treatment at a multi-omics level and basic nematode biology.

## Results

### A comprehensive multi-omics approach to prioritize targets

We have developed a knowledge-driven scoring system for prioritizing nematode intestinal drug target candidates (**Fig. 1A, Fig. 1B**) using the wide range of high-quality genomic, transcriptomic and proteomic data generated from previous studies [19, 21, 23] (and several available annotation programs [24–27]), in the context of phylogenetic relationships, to assure broad control potential. The prioritization (**Fig. 1A**) was applied on the 3,564 *A. suum* genes belonging to Conserved Intestinal Families (cIntFams; conserved expression in *A. suum*, *H. contortus* and *T. suis* [23]), and performed scoring based on orthology, intestinal proteomic evidence [21], high intestinal gene expression (spanning the core species [19, 23, 28], functional annotations including KEGG annotations [29] and RNAi phenotypes in *C. elegans* [30–33], and having many predicted protein-protein interactions [27] (see Methods for additional details). In our preliminary prioritization scheme (**Fig. 1A**), the maximum **gene prioritization score** is 11, and the top-scoring gene (score 10.76) was 2,3-bisphosphoglycerate-independent phosphoglycerate mutase (iPGM, GS_11702). iPGM has previously been identified as a promising macro-filarial drug target for adult filarial worms because it is present, conserved and essential in nematode parasites (and their endosymbiont *Wolbachia* when present) but is absent in humans [34, 35]. Recently, a series of cyclic peptides and analogs exhibiting potent and isozyme-selective inhibition against iPGM orthologs have been developed (“ipglycermides” [36]). The top 50 overall genes and their properties are shown in **S2 Table**.

**Fig. 1.**
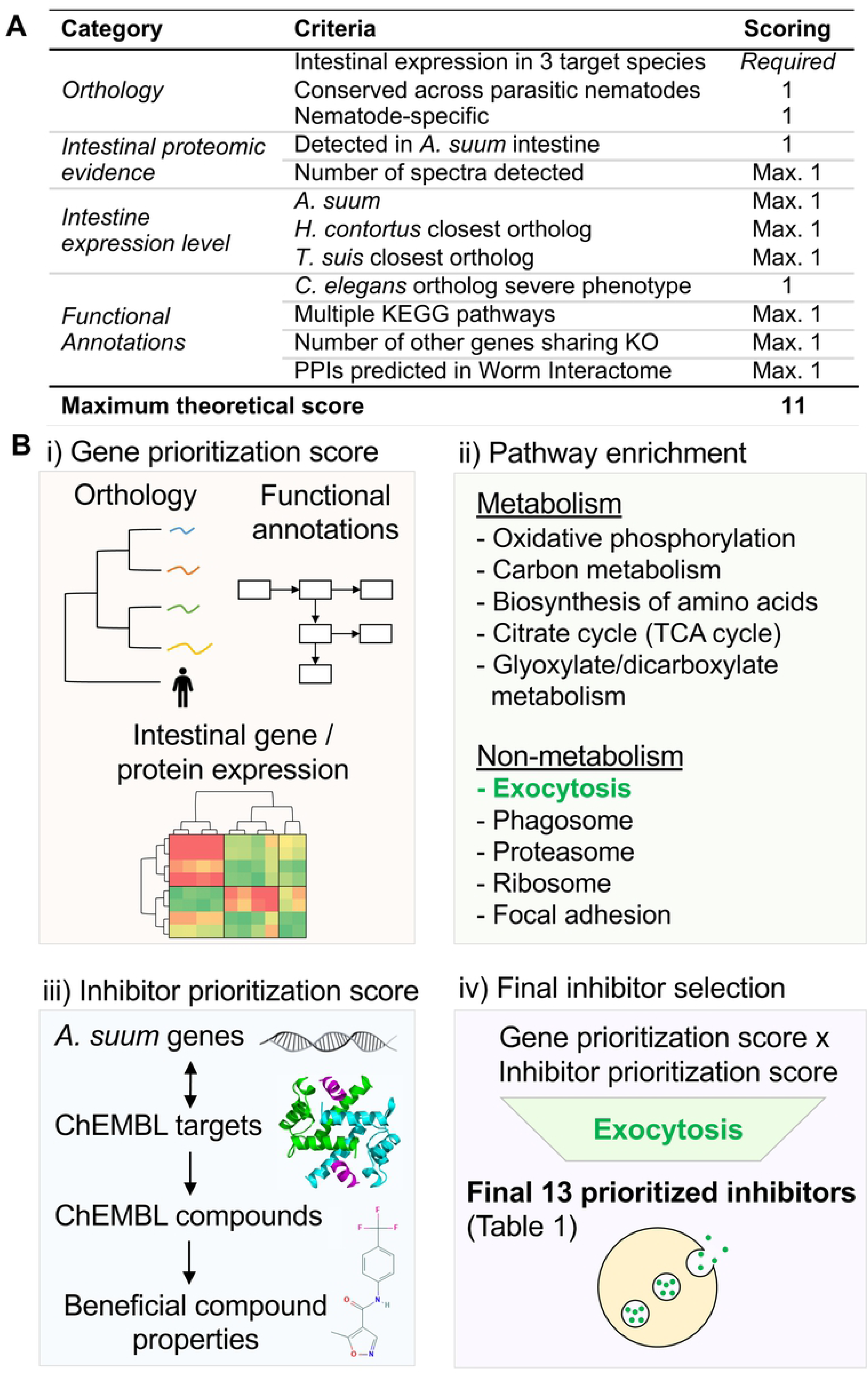
Overview of computational prioritization process. (**A**) Scoring scheme for prioritizing intestinal *A. suum* genes for inhibitor targeting. (**B**) The overall computational target / inhibitor prioritization scheme used to prioritize inhibitors for experimental validation, divided into the broader steps of (i) gene prioritization score calculation, (ii) pathway enrichment testing, (iii) inhibitor prioritization score calculation and (iv) final inhibitor selection. The final 13 inhibitors included 9 computationally prioritized inhibitors and 4 inhibitors manually selected because they are known to target the exocytosis pathway.

### 13 inhibitors are prioritized based on the target prioritization

The overall approach to identify small molecule inhibitors with anthelmintic potential is depicted in **Fig. 1.** Briefly, *A. suum* intestinal genes were first prioritized as potentially good inhibitor targets based on their biological properties (*see* Methods), and based on these results, enriched target KEGG pathways were identified and prioritized. Specifically, “exocytosis/synaptic vesicle cycle pathway was identified as the most significantly enriched. Independently, inhibitors were matched to *A. suum* genes based on homologous targets in the ChEMBL database and scored according to their known properties. The scores from all 3 of these broad approaches were combined to compute a final inhibitor score and to generate a short list of the 25 best-scored inhibitors, expected to target exocytosis related functions. The top 25 inhibitors and their respective scores are shown in **S1 Table**, and included Albendazole (a widely used broad spectrum anthelmintic [37]) as the 16th ranked candidate, which supports confidence in the prioritization approach. Based on availability and cost, a subset of 13 inhibitors was selected for experimental screening using a phenotypic motility assay and parasitic stages of three nematode species spanning the phylum Nematoda. The top 50 gene targets and their detailed scoring criteria are provided in **S2 Table**, all enriched pathways are provided in **S3 Table**, and detailed scoring for all scored exocytosis genes are provided in **S4 Table**.

### Prioritized inhibitors are effective against *A. suum* lung larval stages

Several considerations went into the design of experiments to test inhibitors identified from the screening strategy described above. First to provide for a more inclusive screen, primary assays were done using a relatively high concentration (1 mM) for all inhibitors tested, with the exception of Staurosporine for which a lower starting point (100 µM) was justified by previous findings [38]. Our assessment of movement considered no movement or normal movement by comparisons to controls. We noted slow movement but found these data unnecessary for reaching conclusions on motility assays. Second, although identified with a focus on inhibitors of exocytosis in intestinal cells, inhibitors of this process are likely to impact other cells and other cellular pathways. Therefore, an exclusive impact on inhibition of secretion in intestinal cells is not expected from experiments conducted here. Third, morphologic phenotypes induced by inhibitor treatments are expected to facilitate future identification of anthelmintic targets and dissect underlying mechanisms of potency. Thus, effort was devoted to clarifying whole worm, tissue and cellular phenotypes presented by the experimental system.

#### Effects on motility and molting of *A. suum* L3 identified 10 hits

The 13 inhibitors (**S1 Fig, Fig. 2**) were tested for effects on motility of 8-day old L3 *A. suum* obtained from rabbit lungs (**Fig. 3**). The observed motility inhibition patterns over the period of 5 days could be categorized into four groups: A) relatively rapid acting inhibitors that showed motility inhibition of 70-100% of the worms by 24 hours after treatment (**1**, Leflunomide; **2**, Staurosporine; **3,** Ruxolitinib); B) inhibitors causing motility inhibition of 100% of the L3 after 5 days of treatment (**4**, Combretastatin; **5**, Alvocidib; **6**, Sunitinib and **7**, CID 1067700); C) inhibitors causing motility inhibition in over 70% of the worms but never up to 100% by day 5 (**8**, Taltobulin; **9**, Camptothecin and **10**, Tofacitinib); and D) relatively ineffective inhibitors with greater than 70% of the worms motile (**11**, Podofilox; **12**, KW-2449 and **13**, Fasudil) within the time frame tested.

**Fig. 2:**
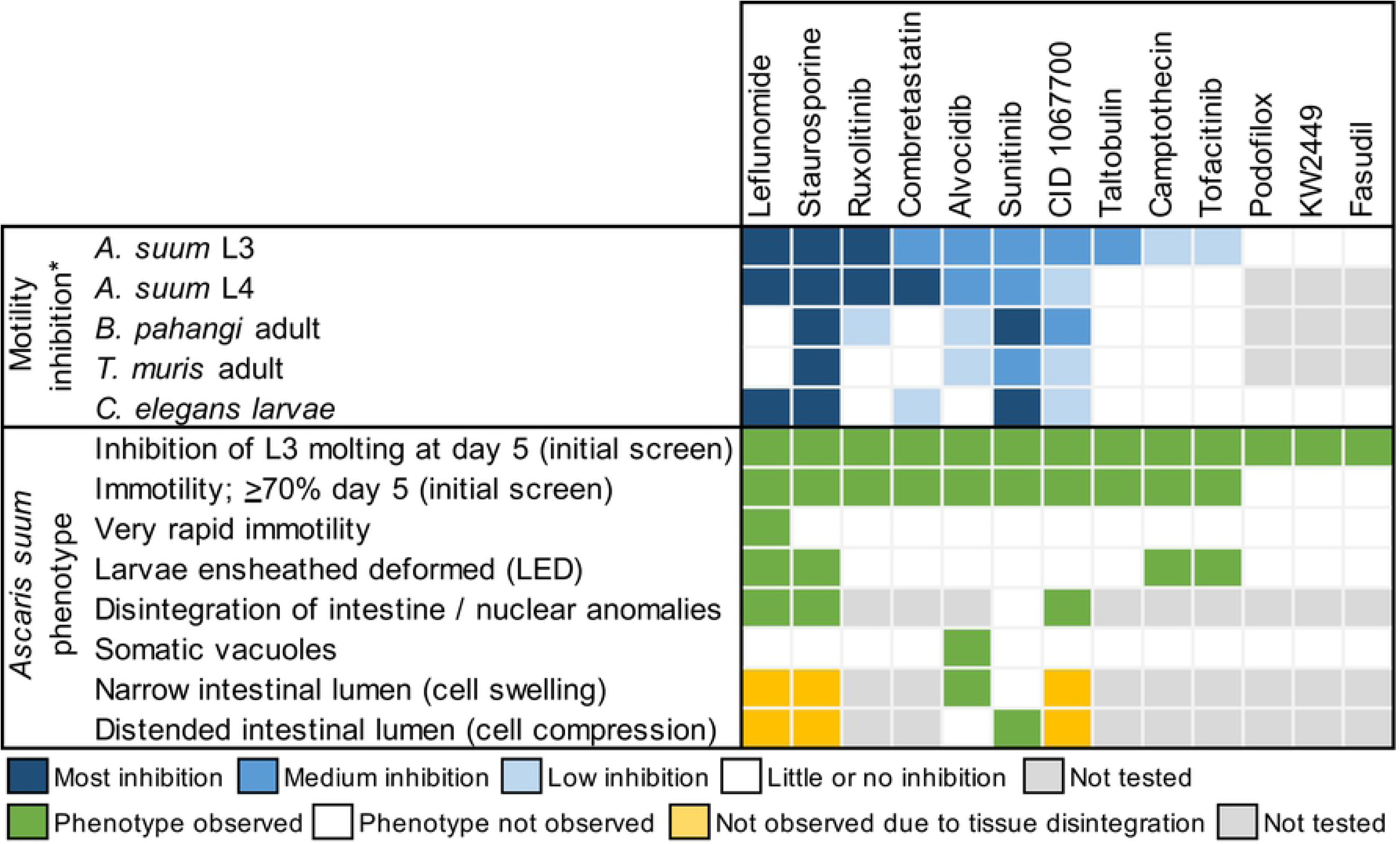
Summary of motility inhibition and observed phenotypes for the 13 tested Inhibitors in four species.

**Fig. 3.**
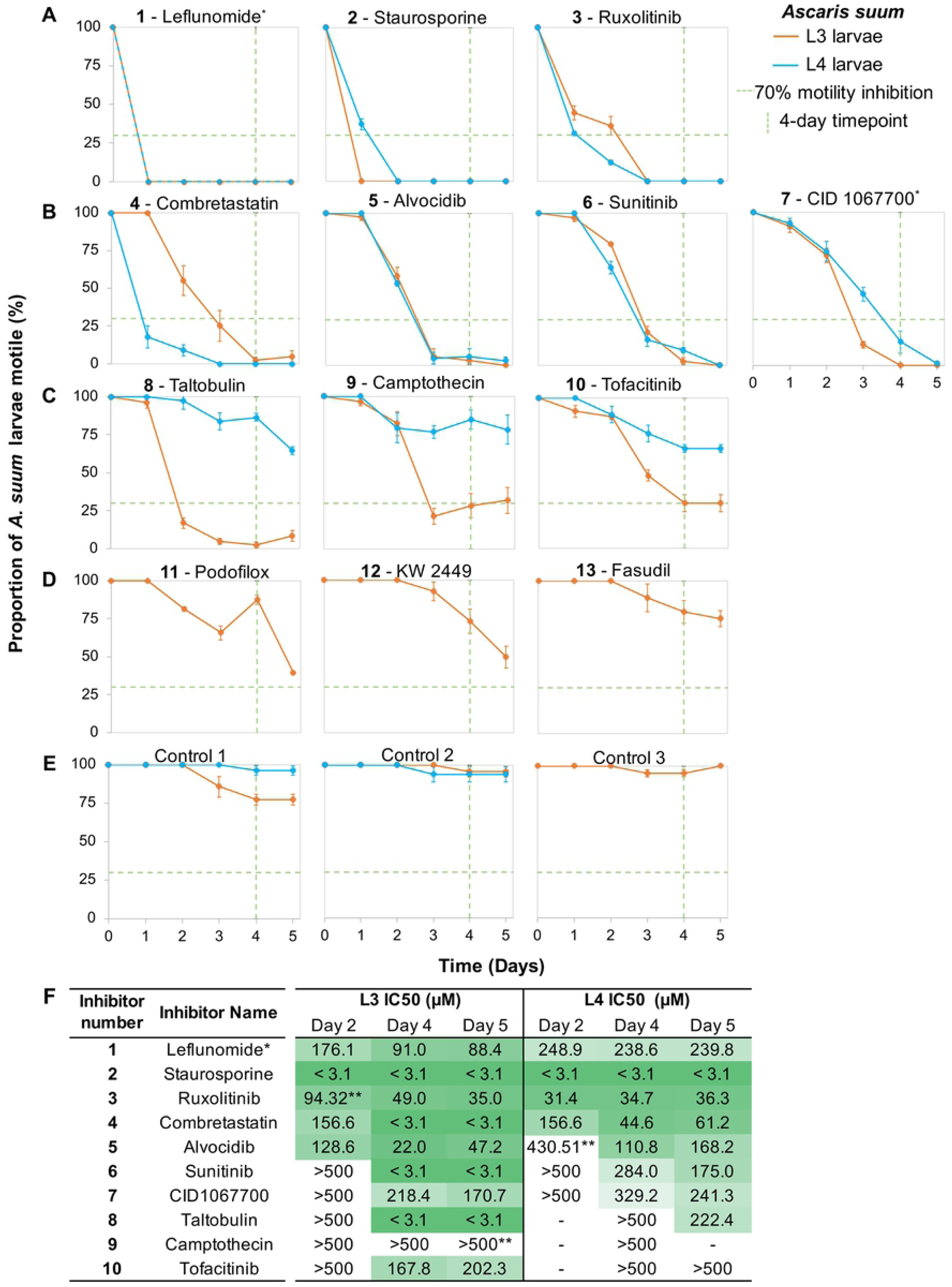
Motility inhibition of inhibitor-treated L3 and L4 *A. suum* larvae (1 mM, except for Staurosporine at 100 μM). (**A**) Inhibitors with rapid and complete efficacy in both L3 and L4 (**B**) Inhibitors with slower but complete efficacy in both L3 and L4. (**C**) Inhibitors with moderate efficacy in L3 and low efficacy in L4. (**D**) Inhibitors with low efficacy in L3 (not tested in L4). (**E**) Controls corresponding to L3 and L4 experiments. Green dashed lines indicate the threshold for inhibitors considered in further testing (≤30% motility inhibition, after 4 days). Error bars represent standard error. *Samples corresponding to Control 1 L3, which had some moderate motility reduction (∼25%) compared to other controls. (**F**) IC_50_ determination for motility inhibition after 2, 4, and 5 days of inhibitor exposure in *Ascaris suum*. *only includes concentrations of 125, 250, and 500µM for L4. **Significant lack of curve fit observed (see **S3 Fig.** for an example of lack of fit).

All 13 inhibitors tested in this primary screen inhibit molting, in that no molting (shed cuticles) was observed in any wells of the treated larvae during the 5-day course of treatments (**S2 Fig.**), whereas shed cuticles occur in wells of control larvae by day 3 in culture (mean 88%, triplicate wells).

The primary phenotypic screens show that each of the 13 inhibitors have some impact on *A. suum* L3, at minimum on molting, with at least 10 that inhibit motility of >70% of worms at day 4 in culture, and hence are considered hits. The 10 hits in L3 were thus tested on *A. suum* L4.

#### Similar effects on motility are observed for *A. suum* L4

*A. suum* L4 experiments focus on the 10 hit inhibitors for L3. The effects on L4s are similar to L3s for 7 inhibitors (**Fig. 3A and 2B**), with inhibitors **1**-**3** inhibiting motility rapidly to the highest levels, inhibitors **4**-**7** having less rapid effects, while **8**-**10** had lower effects on L4 motility which was below the 70% inhibition by day 4 (**Fig. 3C**). Overall 7 of the 10 inhibitors that are effective on L3 also effectively inhibit L4 motility.

#### IC_50_ determinations on *A. suum* L3 and L4

IC_50_ values were determined in motility assays using *A. suum* L3 and L4, although see results below on morphological phenotypes, are more potent on the L3 compared to the L4 on Day 5 of the assays (**Fig. 3F,** see **S3A Fig.** and methods for details). Inhibitor **2** is the most potent of the 10 inhibitors with IC_50_s of less than 1.55 μM after only two days for both L3 and L4, followed by inhibitor **3** with IC_50_s of 94 μM and 32 μM after two days for L3 and L4, respectively), although other inhibitors are comparatively effective after 4 or 5 days (including inhibitor **4**, with IC_50_ of 3.1 μM after 5 days).

Because inhibition of molting reflects a biological process that may have application to targeting by anthelmintics, we also assessed concentrations of inhibitors at which L3 cuticles are first observed in the dilution series. Shed cuticles first occurred at 62.5 μM and below (**1** and **10**), or only at 31.3 μM (**7** and **9**), and no shed cuticles occurred in wells for the remaining 6 inhibitors at any concentration tested. The results indicate that the molting process is quite sensitive to the inhibitors investigated, and apparently as sensitive as, or more sensitive than, overall motility.

### A variety of morphological phenotypes are induced by inhibitor treatments

In many cases, the morphology of immotile worms resulting from inhibitor treatments is either straight or slightly curved as shown in low power, end point (5 days, but achieved earlier in association with lack of immotility) images for *A. suum* L3 (**Fig. 4A**), and representative phenotypes for **2-5**, **7** and **9**. However, some remarkable caveats and exceptions are documented below for **1**, **6**, **9** and **10**.

**Fig. 4.**
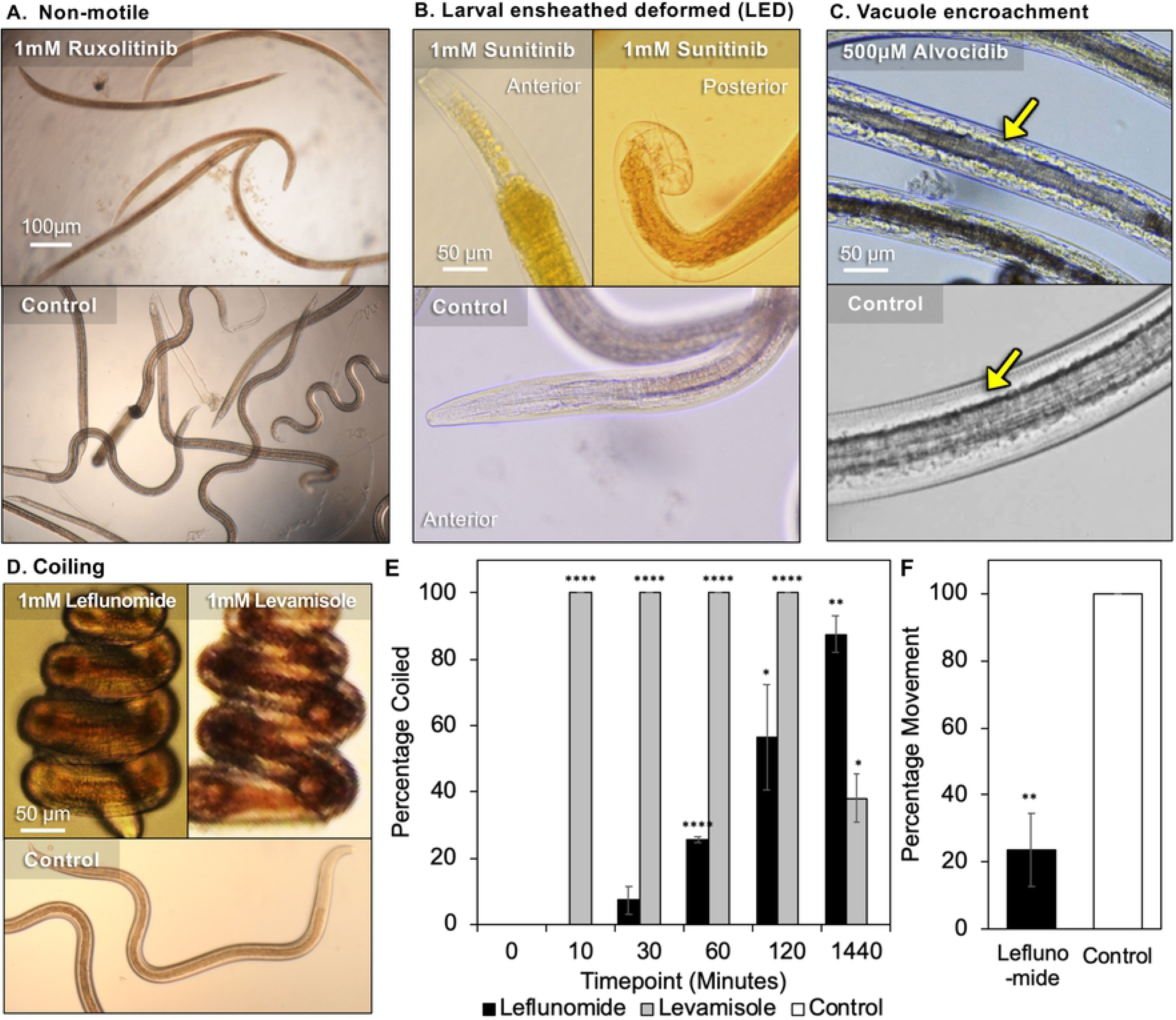
Observed phenotypes among tested inhibitors in *A. suum* L3 larvae. (**A**) Non-motile - Alvocidib, Ruxolitinib, Staurosporine, CID 1067700, Taltobulin, Combretastatin. (**B**) Larvae ensheathed deformed (LED); showing detached cuticle and deformed anterior and posterior ends of the larvae induced by Sunitinib treatment (also, replicated with Tofacitinib, Camptothecin, Leflunomide, Staurosporine). (**C**) Somatic vacuoles (white arrow). (**D**) Coiling (Leflunomide). (**E**) The proportion of larvae with the coiled phenotype over time. Controls were all zero values. (**F**) Motility comparison after 90 minutes of larvae exposure to Leflunomide. For E and F, asterisks represent the results of two-tailed T tests with unequal variance (* P < 0.05, ** P < 0.01, *** P < 0.001, *** P < 0.0001).

Although molting of *A. suum* L3 was inhibited by all 13 inhibitors in the primary screen, an unexpected morphology occurs with 2 inhibitors (**6** and **10**). The morphology involves partial or complete apolysis without ecdysis resulting in larvae ensheathed in the L3 cuticle, and there was obvious deformation of the anterior and posterior ends of worms, leading to the LED phenotype (“larvae <u>e</u>nsheathed with <u>d</u>eformation”; **Fig. 4B**). We interpret this morphology to reflect an abortive molting process from L3 to L4, making it unclear if the effects are on L3, L4, both, or a transition phase between the two. These effects differ from those of the other inhibitors tested that prevent molting from proceeding this far. Hence, the different effects suggest that different inhibitors inhibit L3 at different steps in the process of molting to L4. The unusual LED phenotype that Sunitinib (**6**) and Tofacitinib (**10**) induce includes outright degenerative effects, suggesting activation of degenerative cellular responses that may have value for targeting in anthelmintic strategies.

Inhibitor **5** uniquely induces somatic vacuoles visible within what normally constitutes the pseudocoelomic cavity in L3 and L4 (**Fig. 4C**), and these are obvious by 3 days post-treatment. Additional information on this phenotype is provided below.

Inhibitor **1** induces another whole worm phenotypic effect. **1** routinely Induces a coiled (spiral) morphology in *A. suum* L3 and L4, which also ensues when these larvae were exposed to the neurotoxic anthelmintic levamisole (**Fig. 4D**). However, compared to levamisole, which induces coiling in the population after only 10 minutes, **1** took more than two hours to cause coiling in more than half of the larvae (**Fig. 4E**), although motility is inhibited by almost 80% after only 90 minutes of exposure (**Fig. 4F**). While levamisole treated worms show some recovery after 24 hours, **1** had a longer lasting effect with over 90% of larvae being coiled at 24 hours (**Fig. 4G**). This overall similarity in morphology as induced by levamisole coupled with very rapid immotility suggests that **1** can be neurotoxic in nematodes, a possibility that has not been previously documented for this inhibitor. Nevertheless, **1** also causes extensive cellular and tissue damage, described below, and the final end point morphology with this inhibitor becomes straight to curved, as in **Fig. 4A**, with further time in culture. This second phenotype probably indicates toxicity that extends beyond neurotoxic effects causing the coiled phenotype, which is supported in subsequent experiments described below.

Thus, a variety of phenotypes were documented by gross microscopic assessment that can differentially be attributed to individual inhibitors or subsets of inhibitors tested.

#### Larvae ensheathed with deformation (LED) phenotype in *Ascaris suum* L3

The extensive pathology in association with LED led us to further investigate this phenotype. Both, **6** and **10** are known kinase inhibitors [39, 40] (**S5 Table**), which raises the possibility that inhibition of kinase activity is responsible for inducing LED. However, **2** is a more general kinase inhibitor, but does not noticeably induce LED within constraints of the primary assay. **2** inhibits motility within 24 hours at high concentrations and prevents progression into molting at all concentrations tested. Thus, when presented to L3 at time 0, **2** may act at an earlier step in molting that prevents acquisition of the phenotype induced by the other two kinase inhibitors and masks its ability to induce LED. We tested this hypothesis by adding **2** to L3 after two days in culture and near the time when normal molting occurs. Motility and LED were both assessed on day 5 post-treatment. In this case, the LED phenotype did occur, although some larvae underwent molting, which may reflect variability in the precise developmental position at the time of treatment (not fully synchronized populations; **Fig. 5**). The results indicate that if presented just prior to molting **2** can produce LED, which may indicate that kinase inhibition contributes to causing this phenotype.

**Fig. 5.**
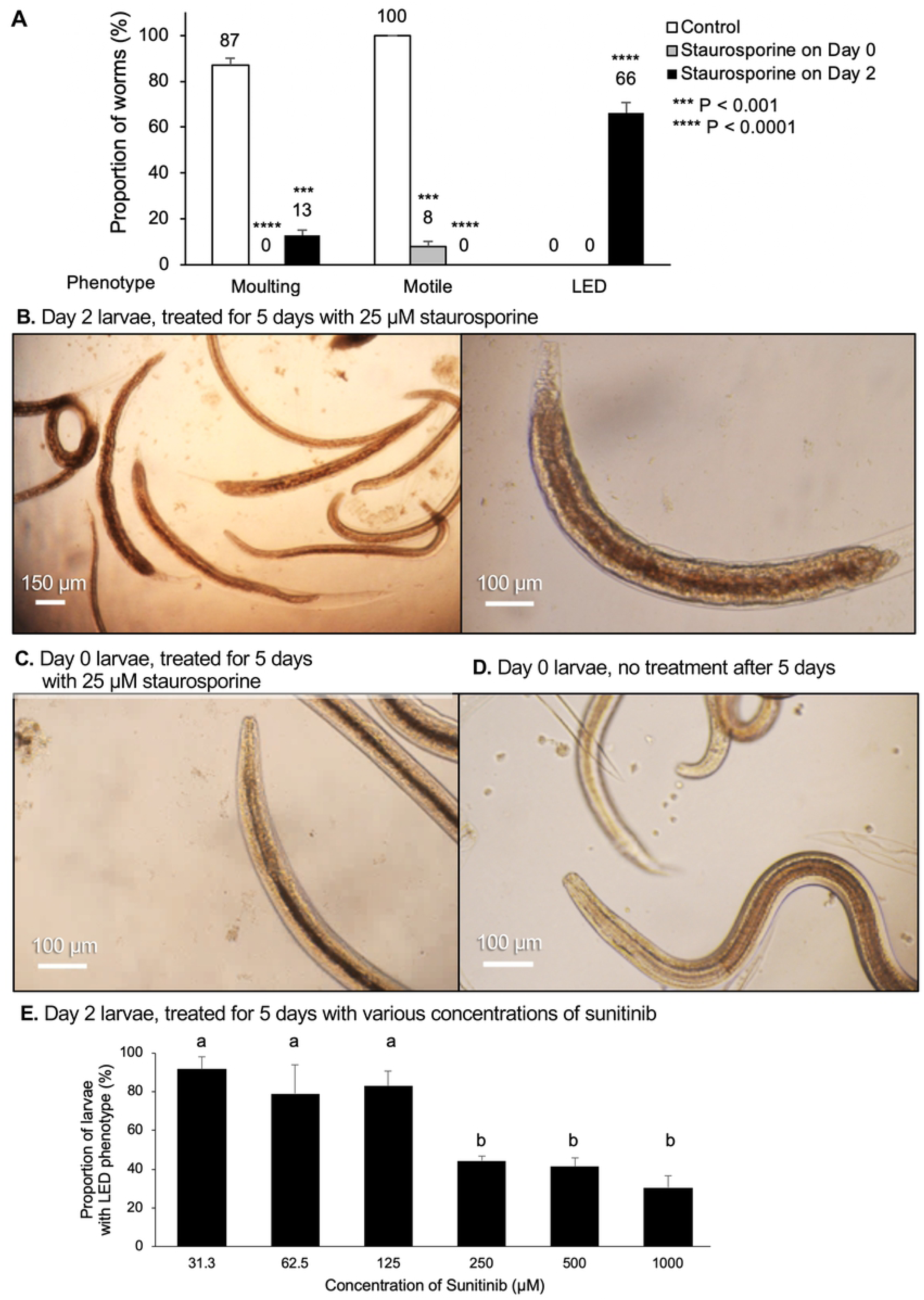
<u>L</u>arvae <u>e</u>nsheathed with <u>d</u>eformation (LED) responses to varying Staurosporine timing and Sunitinib concentrations. (**A**) The observed phenotypes are quantified after 1 µL of 25 µM Staurosporine was delivered to *A. suum* L3 (from 8-day rabbit infections) at time 0 (Day 0) and after 2 days in culture (Day 2), compared to control larvae. P values indicate the results of a two-tailed T-test with unequal variance, compared to control. (**B**) *A. suum* treated at Day 2 demonstrate the non-motile and LED phenotypes, 5 days following treatment. The second panel is magnified to highlight the anterior and posterior damage to the worm. (**C**) *A. suum* treated at Day 0 are immotile and do not have a visible LED phenotype 5 days after treatment. (**D**) Untreated controls are motile and have molted to L4. (**E**) Increasing concentrations of Sunitinib (applied to Day 0 L3 larvae for 5 days) result in a lower occurrence of the LED phenotype. A one-way analysis of variance (ANOVA) indicated significant variation among conditions (*P* = 1.7×10^−5^). Samples grouped significantly into groups a and b (as indicated) according to Tukey HSD *post-hoc* test.

IC_50_ experimental results provided additional insight on the LED phenotype. First, the percentage of larvae that display the LED phenotype increases with decreasing concentrations of **6** over the range used in these experiments (∼30% for 1000 μM, ∼40% for 500 μM and 250 μM, and ∼80% for 125, 62.5 and 31.3 μM; **Fig. 5E**), but molting did not proceed to completion at any concentration used. As with **2**, these results suggest that higher concentrations of **6** inhibit entry, or a very early step, into the molt and prevents acquisition of the LED phenotype. Thus, while lower concentrations of **6** permit increased entry into the molting process, inhibition of a subsequent step(s) appears to lead to the LED phenotype.

A second observation was that rapid inhibition of motility by day 1 post-treatment with **1** is not observed below 250 µM, whereas at 125 µM, the motility curve (**S3B Fig.**) resembles curves observed with **6** and **10** (**Fig. 3B and C**) in that inhibition of motility is delayed and then ensues fairly steeply after 3, or 2 days, respectively in culture. The LED phenotype only occurs at this concentration of **1** in the dilution series, and below 125 µM most L3 remain motile and undergo molting. In addition to DHODH inhibitor effects (**Table 3**), **1** is known to inhibit kinase activity [41]. Thus, the dilution series resolved an additional phenotype that **1** induces and may reflect different levels of inhibition on a single target, or differential sensitivities of multiple targets at different concentrations of **1** in *A. suum* larvae.

Third and last, L3 treated with **9** displayed the LED phenotype at concentrations beginning at 500 µM. Although **9** is a topoisomerase I inhibitor it can also indirectly inhibit kinase activity [42]. In total 5 inhibitors (**1**, **2**, **6, 9, 10**) induce the unusual LED phenotype.

*A. suum* L4 do not progress to L5 in the culture system used here. Consequently, there is no evidence of the LED phenotype in L4s at any concentration of the top 10 inhibitors tested. From a morphologic perspective, the analysis is complicated by the fact that less than 100% of L3 molt to L4 (mean 88%, triplicate wells), and hence the remaining L3 which do not undergo molting might still be able to acquire the phenotype, if induced. Indeed, a phenotype resembling LED sporadically occurs in experiments involving L4 larvae. We attribute this low-level occurrence of the phenotype to the presence of unmolted L3.

### Inhibitor effects are observed in phylogenetically diverse nematode species

*A. suum* is a clade III representative of the Nematoda[43]. The 10 inhibitors that are hits in *A. suum* L3 and L4 (hit defined as having motility inhibition of at least 70% by day 4; **Fig. 3**) were next tested *in vitro* with adult *B. pahangi* (another clade III representative) and adult *T. muris* (a clade I representative) (**Fig. 6**). Single dose preliminary screens were carried out at 100 µM for all except **2** (25 µM). Using the same motility cut-off of at least 70% inhibition but over 6-day exposure (due to 10-fold lower concentration used compared to the primary *A. suum* screen), 6 inhibitors are hits in *B. pahangi* (**2**, **3, 5**-**8**), although response to **3** is variable between days 5 and 6. Interestingly, 5 of these inhibitors (**2, 3, 5, 6** and **7**) are also effective in inhibiting motility of adult *T. muris*. Thus, each of these 5 inhibitors (**2, 3, 5, 6** and **7**) are hits with all 3 parasitic nematode species representing nematode clades I and III, whereas **8** is a hit for *A. suum* and *B. pahangi*, but not *T. muris*, leaving 4 additional inhibitors as hits (**1, 4, 9,** and **10**) only for *A. suum* larvae.

**Fig. 6.**
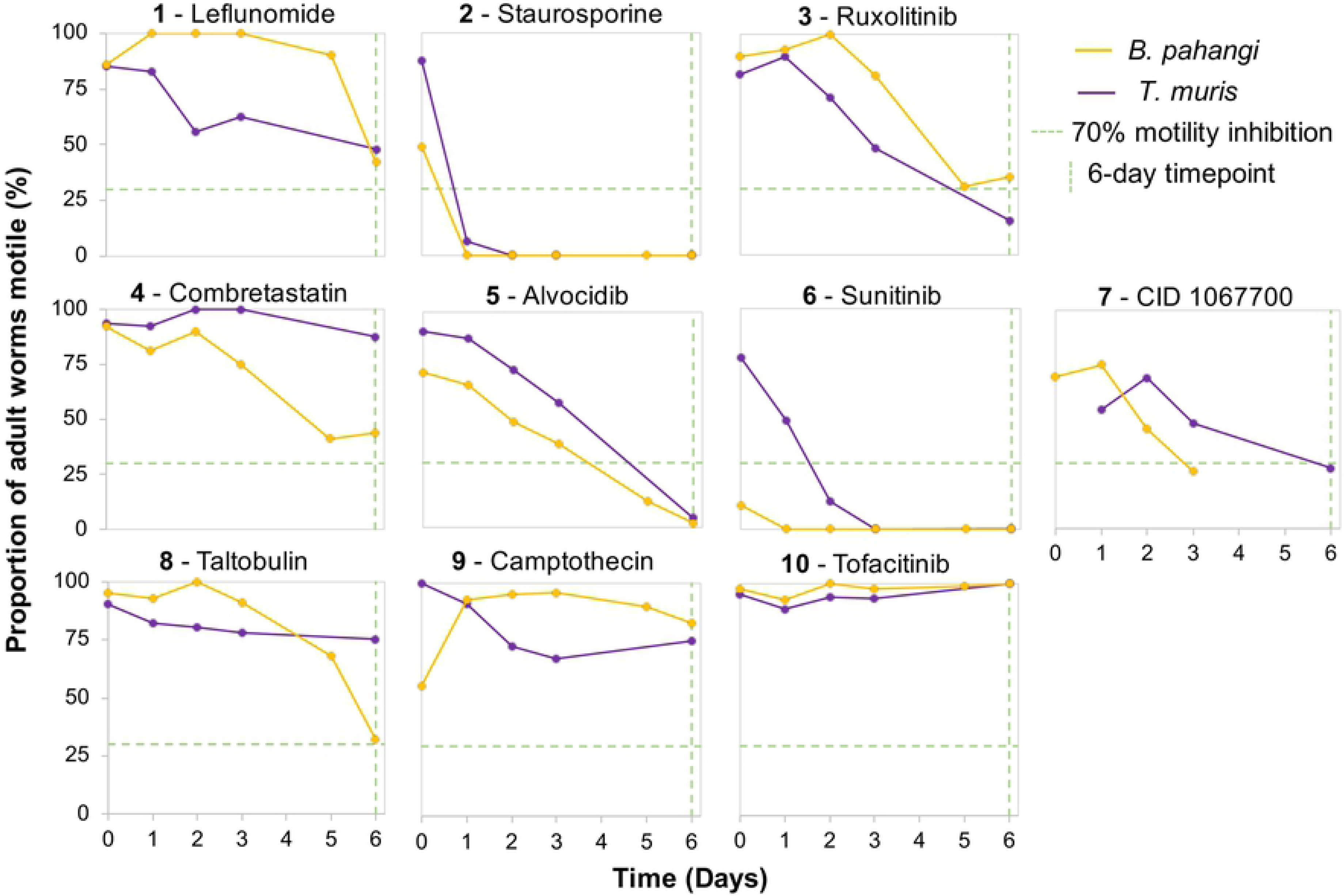
Motility responses of inhibitor treated *B. pahangi* and *T. muris* (100 µM, except for Staurosporine at 25 µM).

Four of the top 10 inhibitors that were tested with *A. suum* L3 (**2**, **6**, **9** and **10**) were tested with *B. pahangi* L3. As all 4 of these inhibitors interfered with motility and molting at some concentration, they were selected primarily on the basis of their ability to induce the LED phenotype in *A. suum* L3. **2** at 25 µM inhibits motility by 73% within 24 hours following addition of the drug and inhibits larval motility by 100% on day 7, the day before most control larvae begin to molt. Thus, **2** apparently kills the L3 before they can molt. Sunitinib (**6**) at 8 µM, on the other hand, inhibits larval motility by 44% but completely inhibits larval molting on day 8 suggesting that even when larvae are actively moving, they are unable to continue through the molting process. Consequently, both **2** and **6** had similar effects on *A. suum* L3, i.e. these drugs inhibit both motility and molting. Interestingly, **9**, a topoisomerase 1 inhibitor, does not appear to inhibit motility throughout the course of the 12-day assay but it does inhibit molting by 79% with 62.5 µM by Day 12. **10**, which is a JAK1 and JAK3 inhibitor, does not appear to have a significant effect on larval motility nor molting even at a higher concentration of 125 µM. By contrast to *A. suum*, none of the inhibitors cause a phenotype resembling LED in *B. pahangi* L3. There was an occasional lesion evident emanating from the body of L3 treated with **6**, by comparison to controls (**S4 Fig.**). Nevertheless, *B. pahangi* L3 that fail to molt, even in untreated cultures occasionally exhibit a phenotype resembling LED (**S4 Fig.**), and a related phenotype has been more commonly observed in cultures of *B. pahangi* L3 lacking exogenous ascorbic acid (vitamin C). Thus, while LED is potentially inducible in *B. pahangi* L3 the conditions required to initiate it may differ among nematode species.

*C. elegans* is a clade V nematode representative and as a well-developed model nematode offers potential to investigate mechanisms and cellular targets by which inhibitors confer toxicity. Each of the original 13 inhibitors were also tested in motility assays with *C. elegans* initiated with 1 day or 2 days old larval stages (**S5 Fig.**). Of the 13 inhibitors tested, three (**1**, **2**, **6**) inhibit motility of over 70% for both 1 day and 2 days old larvae by 2 days of treatment. Several other inhibitors cause moderate levels of inhibition (20% to 70%) for 1 day and/or 2 days old larvae (**3**, **4**, **7**-**9**), while several have minimal (<20%) to no effects on either (**5**, **10**, **12**, **13**). Thus, inhibitors **2** and **6** have activity based on our definition of ‘hit’ in each of the 4 species tested, while **1** is a hit for both *C. elegans* and *A. suum*. Next the ability of inhibitor **1** to very rapidly inhibit motility in *C. elegans* larvae was tested in 30 minute assays, and it was found to do so (**S5B Fig.**). In this case, larvae do not present with a coiled phenotype as in *A. suum*. Regarding the LED phenotype and in contrast to *A. suum*, **6** does not appear to cause this phenotype in *C. elegans* in assays at 1 mM, although it does cause high level immotility within 24 hours. Nevertheless, no larvae present with the LED phenotype in assays using lower concentrations of **6**, or with other chemical inhibitors tested against *C. elegans*. Overall, *C. elegans* responses indicate potential to aid in dissecting anthelmintic mechanisms for some, but not all of the inhibitors that displayed activity against the parasitic species tested.

### Prioritized inhibitors induce an array of intestinal cell and tissue pathology

To investigate the hypothesis that inhibitors identified so far are toxic to intestinal cells, we established three complementary approaches that were applied to *A. suum* L3: 1) use of DIC microscopy, 2) live worm bisbenzimide nuclear staining, and 3) standard histological staining of sections of treated worms. Experiments were focused on *A. suum* L3 treated with each of 5 inhibitors (**1, 2**, **5**, **6, 7**) that were selected based on overall performance in collective experiments and diversity of potential targets. Concentrations were chosen that were likely to cause pathology by day 2 of treatment (25 µM for **2**; 500 µM for all others). Assessment for DIC and bisbenzimide assays focus on the region immediately posterior to the intestino-esophageal junction to provide consistency across treatments. This localization was not possible for tissue sections because the small size of larvae precluded an ordered orientation in the histological preparations.

DIC microscopy resolved general tissue and cellular characteristics of control larvae, to the extent that intestinal cells show apparent outlines of cell membranes and nuclei (**Fig. 7A**). In contrast, larvae treated with **1**, **2**, **7** show vacuolization and otherwise disruption of intestinal cell organization (**Fig. 7B, 6C, 6F**), which is relatively extreme with **7** in that no cellular organization is evident between the basal margins of the intestine (**Fig. 7F**). L3 treated with **5** and **6** appear to have a more normal pattern for intestinal tissue (**Fig. 7D, 6E**), although the yellow background stain of **6** interferes with resolution of effects by DIC (**Fig. 7E**).

**Fig. 7.**
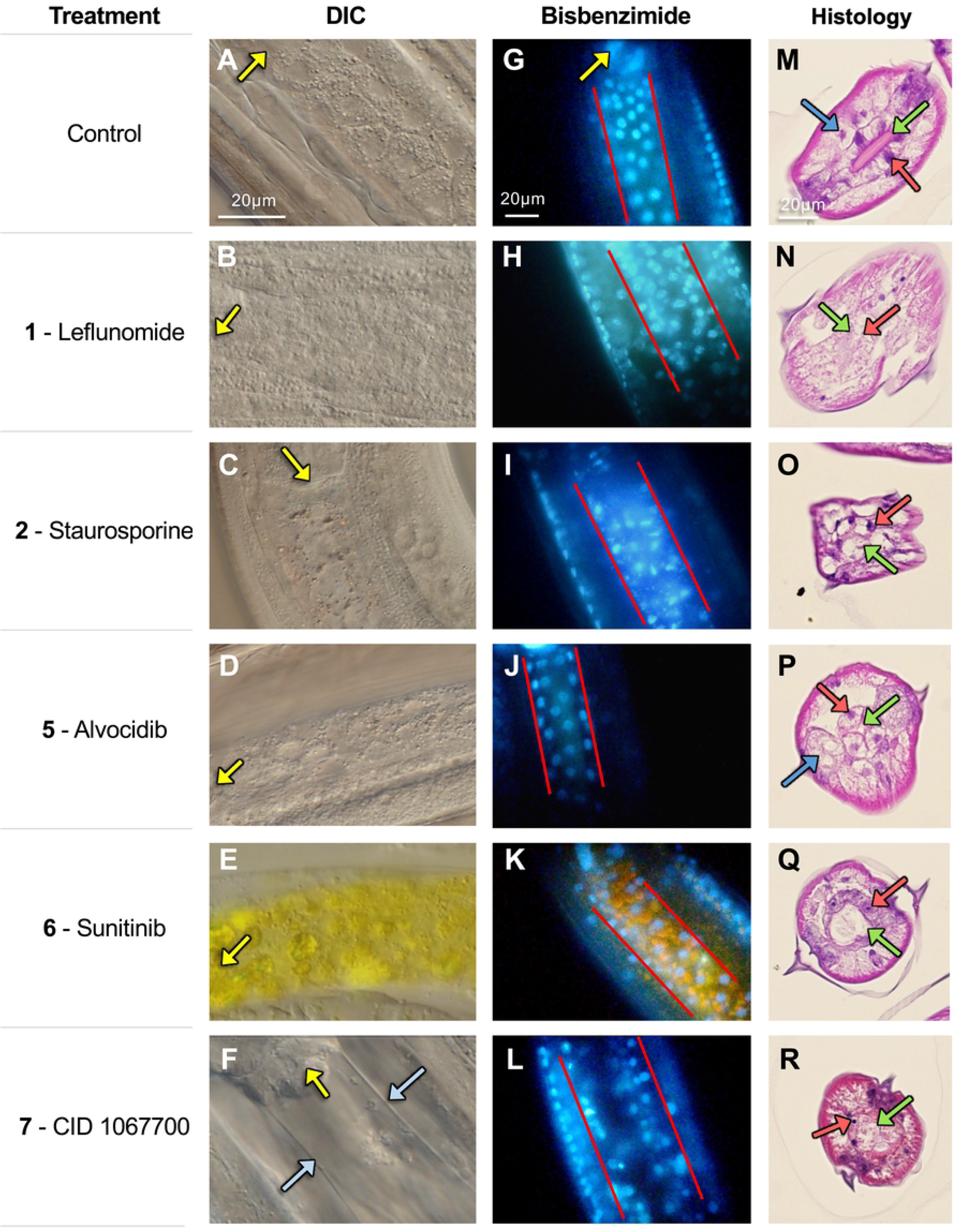
Microscopy of L3-stage *A. suum* treated with five of the prioritized inhibitor candidates (500 μM, except Staurosporine at 25 μM) at day 2. (**A-F**) Differential interference contrast (DIC) microscopy, with the pharynx positions indicated by yellow arrows and the intestinal phenotype for CID 1067700 indicated by blue arrows. Note that Sunitinib itself fluoresces yellow. (**G-L**) Bisbenzimide fluorescent staining indicating the nuclei and nuclear morphology in treated larvae. The orientation of the pharynx is indicated in the control image with a yellow arrow. Red lines indicate the boundaries of the intestine. (**M-R**) Histopathology of larval cross-sections stained with eosin and hematoxylin. Red arrows point to intestinal cell nuclei, green arrows point to the intestinal lumen and apical intestinal membrane, and blue arrows indicate the enlarged muscle cells resulting from Alvocidib treatment. In some cases, the arrows point to apparent locations of these structures because of the level of histopathic damage. Images shown are representative of five biological replicate images taken for each inhibitor (with no substantial variability among replicates for any of them).

Following treatment with **5** and **6** in bisbenzimide assays, L3 intestinal cell nuclei routinely show fully rounded morphology and regular distribution of nuclei similar to that with control L3 (**Fig. 7J, 7K, 7G**, respectively). In contrast, treatments with **1**, **2** and **7** each induce altered nuclear morphology and disruption of the regular distribution of intestinal cell nuclei within tissue (**Fig. 7H, 7I, 7L**) compared to control L3 (**Fig 7G**), suggesting significant histological damage to intestinal cells and tissues. Histopathologic comparisons confirm this suggestion and demonstrate a loss of intestinal tissue integrity and either pyknotic or poorly stained nuclei in intestinal cells of L3 from these treatments (**Fig. 7N, 7O, 7R**) by comparison to control L3 (**Fig. 7M**).

L3 treated with **5** and **6** also display histopathologic changes by comparison to control L3, but not the gross cellular degeneration observed with the other inhibitors. With **5**, a noticeable reduction in the diameter of the intestinal lumen occurs, and intestinal cells show enlargement (swelling), along with a similar change involving swelling of muscle cells (**Fig. 7P**), as was independently observed in end-point morphology (**Fig. 4C**). L3s treated with **6** regularly display less organized apical membranes, as indicated by altered and less defined staining at the apical surface, a distended and more rounded lumen, and more compressed intestinal cells (**Fig. 7Q**), with some variable presentations of nuclei not evident in the bisbenzimide assay. Although information from other tissue assays do not explain the **6**-induced alterations to the apical membrane (**Fig. 7Q**), the distended lumen of **6**-treated worms may reflect less turgor pressure of intestinal cells, leading to less volume and enlargement of the lumen compared to control L3. The overall presentation with **6**-treated larvae is opposite to the swollen appearance of intestinal and muscle cells associated with **5-**treated larvae (**Fig. 7P**).

Collectively, the data support the hypothesis that the selection process can identify inhibitors that are toxic to *A. suum* intestinal cells. At least 5 inhibitors selected by our model confer toxicity to intestinal cells of *A. suum* L3. In addition to intestinal tissue, treatments of L3 with the inhibitors cause apparent pathology to other tissues, including what appears to be frank disruption of tissue, pyknotic nuclei in lateral lines (hypodermal and apparent seam cell nuclei) among other morphologic changes (**Fig. 7N-6R**), which will be more fully documented elsewhere.

At this point, an array of different phenotypic effects, summarized in **Fig. 2**, reflect pathologic signatures that distinguish among the 13 inhibitors investigated in this research. Although the selection process was guided by a bias to elements of exocytosis, potential to inhibit functions of multiple proteins was integrated into the selection process, and the different signatures support that individual inhibitors impact distinct cellular targets that collectively confer an array of distinct pathologic sequelae, specific to individual or subgroups of inhibitors. Because many pathways either directly or peripherally converge on exocytosis, the possibility of this outcome was anticipated and adds to the significance of our findings. Because of the irreparable damage these inhibitors showed to the nematode intestinal cells we classified them as nematode intestinal toxicants.

## Discussion

Research reported here integrates information from a multi-omic, pan-Nematoda approach focused on investigating the biology of a single tissue of parasitic nematodes, the intestine, to derive cellular pathways and predict targets for which existing approved drugs (or prospective inhibitors) have potential pharmacologic applications. Information distilled from this approach identified exocytosis as a compelling pathway for investigation of the nematode intestine. Information on existing drugs and drug targets in the ChEMBL database was interleaved with parasite intestinal cell information to select inhibitors for testing. A system utilizing *A. suum* lung stage larvae [44] was used for experiments and improved upon by incorporating methods that facilitate demonstration of pharmacological effects on intestinal cells. In all, 13 of 13 inhibitors selected and tested (either approved drugs or rationally selected inhibitors) inhibit at least one process (molting), the lowest effective concentration of which varies among inhibitors. 10 inhibitors meet threshold levels of inhibition of motility for *A. suum larvae* (≥70% mean inhibition by day 4 post-treatment when used at 1 mM (100 μM, Staurosporine), and 5 out of 5 of these selected for further investigation cause demonstrable intestinal cell/tissue pathology in *A. suum* L3 (at 500 μM, except Staurosporine, 25 μM). An overlapping subset of 5 inhibitors cause pathology in context of the LED phenotype. Importantly of those tested, several of the inhibitors score as hits against the adult (5 inhibitors) and L3 (3 inhibitors) of the filarial nematode *B. pahangi*, the adult whipworm *T. muris* (5 inhibitors), and larvae of the non-parasitic nematode *C. elegans* (one or more larval stages, 4 inhibitors). Thus, several of the inhibitors identified have activity against nematode representatives from clades that span much of the phylogenetic diversity of the phylum Nematoda. The predicted specificities of the inhibitors tested, the nature and diversity of pathologic sequelae that they induce, and specific endpoint data from assays established in this research will collectively inform ongoing research and provide methods to investigate underlying mechanisms and cellular targets responsible for the anthelmintic effects observed.

### Evidence-based computational target and inhibitor prioritization

The computational scoring procedure (**Fig. 2**) utilized existing genomic, transcriptomic and proteomic datasets to strategically select protein targets to be matched to inhibitors, based on existing knowledge. Similar approach at a smaller scale or aimed at specific processes has been successfully applied before (e.g. [38, 45]), but in this case our focus was on a single tissue, highlighting this computational approach’s flexibility when studying specific tissues or pathogens or other systems of interest. The prioritization includes criteria to select targets based on orthology (conserved across parasitic nematode species), intestinal proteomic detection (providing evidence that the gene target is active and produces proteins in the worm intestine), high gene transcription levels in the intestine across several nematode species (providing supporting evidence of cross-species conserved intestinal function), and knowledge-based evidence of the gene’s biological functions (providing evidence that inhibiting the target could be lethal in the nematodes based on RNAi phenotype of orthologous targets in *C. elegans*). The prioritization procedure identifies Albendazole (a widely used broad spectrum anthelmintic [37]) as the 16th ranked candidate, which adds confidence for the rest of the prioritized list. Although unclear that inhibitor hits identified will have practical anthelmintic applications, several interesting and diverse pathologic phenotypes were induced, rendering both the actual cellular targets and the mechanisms involved of interest to elucidate. Nevertheless, several inhibitors are approved drugs or in clinical trials (**S5 Table**) for use in humans, which can hasten achievement of anthelmintic applications if warranted. Therefore, the approach was unusually successful in producing a rich source of results from which future research can be prioritized. It seems clear that both refinement and extension of the general approach has applications toward further dissecting basic functions of parasitic nematodes that are essential for their survival. Future bioinformatics prioritizations with different goals can utilize the database constructed for this study or can be built with similar criteria for phylogenetic conservation, functional annotations, evidence of expression, and favorable matched inhibitor properties.

### Link to exocytosis

Despite factoring in exocytosis databases for selection of inhibitors, demonstrating inhibition of exocytosis or secretion in intestinal cells has been challenging and is complicated by many technical hurdles that will require research effort to overcome. This consideration notwithstanding, at least 5 selected inhibitors produce detectable pathology in intestinal cells of *A. suum* L3, 3 of which (Leflunomide, Staurosporine and CID 1067700) caused outright cellular disintegration, nuclear anomalies and nuclear disorganization within intestinal tissue. The other 2 (Alvocidib and Sunitinib) cause morphologic changes (narrow lumen and swelling of cells, or distended lumen and compression of cells, respectively), but without the cellular disintegration regularly observed with the first 3. Both Alvocidib and Sunitinib also induce other pathologic changes discussed below. Therefore, while not demonstrating a direct link with exocytosis, our approach identified inhibitors that are highly toxic to intestinal cells. This general outcome was anticipated and made the approach an attractive one to investigate and potentially identify novel therapeutics despite the challenges of providing direct links to exocytosis.

### Pathologic effects of the active inhibitors

The array of phenotypic changes identified (summarized in **Fig. 2**) go far beyond simple inhibition of movement or molting. These more specific pathologic effects represent processes that when disrupted have lethal consequences for the parasite, and the range of different pathologic presentations must involve diverse targets and cellular pathways. As such, the pathologic presentations provide criteria to begin to uncover the different mechanisms involved. The more remarkable presentations include the LED phenotype (Sunitinib and others), the very rapid immotility response (Leflunomide), and frank disintegration of intestinal cells and tissues (Leflunomide, Staurosporine and CID 1067700).

#### LED

The LED phenotype involves the process of molting from L3 to L4, with apolysis but not ecdysis, followed by deformation (gross disintegration) of larval tissue most obviously at the head and tail of the presumptive *A. suum* L4. This unusual phenotype is distinct from one in which inhibition either prevents entry into molt or blocks earlier steps prior to apolysis, which was induced by all inhibitors tested at concentrations used in the initial screen. Inhibition of molting by protease-class specific inhibitors was previously reported for *A. suum* L3 [46], but, with no reference to an LED-like phenotype. The tissue disintegration with LED is also distinct from well-established examples in which apolysis but no ecdysis occurs, with naturally ensheathed infective L3 of the Strongylid nematode pathogens, as one example, that infect humans and animals, or in *C. elegans* molting mutants [47, 48]. Further, realization of the full phenotype apparently depends on progression of L3 into a susceptible phase of the molting process. For instance, Sunitinib either induced frank immotility of unmolted L3, or the LED pathology, while decreasing concentrations in our dilution series unexpectedly elevate the occurrence of LED, and no minimal concentration was discerned in this series. Further, addition of Staurosporine at day 0 of L3 culture caused rapid immotility and no LED, but when added at the interface of the molting process (day 2 of L3 culture), LED became evident. Thus, toxic effects conferred prior to some specific molting step apparently can prevent induction of the LED phenotype by inhibitors unless specific concentration or timing requirements are met.

All five inhibitors (Leflunomide, Staurosporine, Sunitinib, Camptothecin and Tofacitinib) found to induce LED can be linked to kinase inhibition, which may indicate involvement of kinase inhibition in inducing the phenotype. Nevertheless, testing of a wider range of inhibitors with different specificities at day 2 of L3 culture could be informative here. In any case, LED-inducing inhibitors have identified a distinct developmental step(s) later in molting that when inhibited prevents ecdysis and activates a destructive pathologic response at the L3-L4 developmental interface in *A. suum* larvae. Because of the remarkable destruction associated with this phenotype, the mechanism of activation and mediators of this pathologic process are of interest to elucidate. One possible lead relates to the loss of volume evident in intestinal cells of *A. suum* L3 following treatment with Sunitinib, suggesting dysfunction of cellular fluid regulation. Hence, while not necessarily linked to intestinal cells, the LED phenotype could stem from other cells experiencing similar toxicity. In contrast, inhibitors that induce LED in *A. suum* larvae fail to do so with *B. pahangi* L3, although an LED-like phenotype occurs occasionally with these larvae that fail to molt completely, even in control wells as with *A. suum*, raising the possibility that the phenotype might be inducible in *B. pahangi* under yet identified permissive conditions.

The unexpected findings on LED coupled with the inhibition of entry into molting caused by all inhibitors tested on *A. suum* has potential practical value given that larval stages of numerous nematode pathogens are targets in existing control strategies, e.g. hypobiotic larvae of many strongylid nematodes (including hookworms [49]), and vector transmitted L3 of filarial nematodes such as the heartworm in the vertebrate host [50], as some examples. Relatedly, it may be significant that inhibitors such as Staurosporine and Sunitinib each blocked molting of *B. malayi* from L3 to L4 and caused immotility of adult worms. The pathologic relationship of these observations with those from *A. suum* larvae, inclusive of intestinal cell pathology (**Fig. 7**) is yet unclear and our observations make this a topic of research interest.

#### Very Rapid immotility

In addition to inducing LED at a narrow concentration range, Leflunomide was unique among inhibitors tested in very rapidly causing immotility after exposure of *A. suum* L3 and L4. Affected larvae display a tightly coiled morphology in high percentage that occurs following treatment with the neurotoxic anthelmintic levamisole. A related morphology sporadically occurs with other treatments and even in untreated *A. suum* larvae, but not to the level induced by treatment with Leflunomide (see **Fig. 4E**). *C. elegans* larvae also are very rapidly immobilized by Leflunomide, but do not display the coiled morphology, as is the case for levamisole treatment of *C. elegans*. Thus, Leflunomide appears to have neurotoxic effects that have not previously been reported for this drug in nematodes. Toxic effects of Leflunomide on *C. elegans* were reported [51], but observations were made only after 12 hours, obviating detection of more immediate effects. Leflunomide is an approved drug for treatment of rheumatoid arthritis and its principle mode of action is inhibition of DHODH and mitochondria-based synthesis of pyrimidines, one of two pathways that typically supply pyrimidines to cells. Reversible neuropathy has been noted following treatment with this drug in human patients [52]. Nevertheless, Leflunomide can inhibit other cellular targets, including kinases [41], and it induces several diverse effects in *A. suum* larvae (very rapid immotility, LED, disintegration of intestinal cells). Whether the diverse effects relate to a focal point of disruption expressed differently among tissues or involve disruption of diverse targets and pathways remains to be determined. What is clear is that Leflunomide causes multiple phenotypes that have potential value in application to anthelmintic approaches and warrants further investigation. Findings that *C. elegans* displays at least one of the phenotypes, rapid immotility, may identify a direction that can be taken to investigate this effect.

#### Disintegration of intestinal cells

Separate from the preceding pathologic outcomes, remarkable intestinal cell and tissue disruption ensues in *A. suum* L3 with treatment by 3 inhibitors (Leflunomide, Staurosporine and CID 1067700). For each, the normal regular cell morphology observed by DIC microscopy is disrupted, the normal morphology and distribution of intestinal cell nuclei become altered as shown in bisbenzimide assays, and intestinal tissue displays disintegration along with altered morphology and staining of nuclei in histopathologic sections. Although evaluated with 5 of the original 13 inhibitors, not all that were tested induce this pathologic profile (e.g. Alvocidib and Sunitinib) under the experimental conditions used. Even though all 5 inhibit L3 molting at the concentrations used, the additional pathologic presentations characteristic of each indicate specificity according to the inhibitor, and hence specificity of the target(s)/pathway(s) they disrupt, and the pathologic mechanism(s) involved. The altered distribution of nuclei in the bisbenzimide assays suggests disruption of cell membranes, which is supported by histopathologic results for each of the 3 inhibitors under discussion. Altered shapes of nuclei observed in bisbenzimide assays do not address DNA content, which may be reduced, and both pyknotic nuclei and poorly staining nuclei are apparent on histopathologic analysis of tissue sections. Thus, while not identical, the general pictures agreed between these two complementary methods for each of the 3 inhibitors, and each complementary method adds to more general information provided by DIC microscopy.

Elements of cellular processes leading to the disruption of intestinal cells are not clear from our results, only that the three different inhibitors induce similar features of pathology. CID1067700 is a reported Rab GTPase inhibitor, and Rab GTPases evidently are key regulators of endocytosis, which ultimately influences exocytosis in *C. elegans* intestinal cells [53, 54]. In addition to DHODH, Leflunomide can inhibit kinases (PTK2B) [55] and is an agonist for aryl hydrocarbon receptor (AHR) [56], and it inhibits secretion in inflammatory cells [57] by yet unresolved mechanisms. As a more general inhibitor of kinases, Staurosporine has potential to inhibit a wide array of intestinal cell functions, inclusive of exocytosis and others. Thus, while inhibition of exocytosis is a possible antecedent of the pathology described, there are multiple other possibilities which may vary according to inhibitor. More importantly at this point, it is clear that pathological processes can be induced by diverse inhibitors in intestinal cells of *A. suum* larvae, apparently reflecting irreparable damage. Again in this case, the mechanisms of induction and mediators of damage are of keen interest to elucidate, as they each may represent high value targets for anthelmintics. Of possible relevance here, necrotic processes can be induced in intestinal cells of *C. elegans* by multiple different stimuli, and apparent protease mediators of this pathology can vary according to the stimulus applied [58]. As found here, toxicity of two of the inhibitors, Leflunomide [51] and Staurosporine [59], was shown for *C. elegans*, thus identifying *C. elegans* as a possible resource to dissect mechanisms of intestinal pathology induced by these inhibitors in both species.

### Microtubule inhibitors

Because of the pathology induced by benzimidazole anthelmintics in intestinal cells of parasitic nematodes, we selected two additional inhibitors that bind beta-tubulin (Combretastatin and Taltobulin) for our experiments, and Podofilox was identified by the screening process. Albendazole is an effective anthelmintic and had an IC_50_ of about 3.8 mM *in vitro* motility experiments with *A. suum* L4 (isolated from swine [60]). Although not strictly comparable, Combretastatin and Taltobulin each caused immotility of *A. suum* L3 at 1 mM (IC_50_s of 3.1 and 2.1 µM at day 5, respectively), while Podofilox had more modest effects on L3 motility. Combretastatin was significantly more effective in inhibiting motility of *A. suum* L4 than Taltobulin (IC_50_s 61.2 and 222.4 µM at day 5, respectively). Similar to benzimidazoles, Combretastatin binds at or near the colchicine domain of beta-tubulin [61], whereas Taltobulin binds at or near the vinca domain [62], together providing some diversity in coverage of tubulin domains. Although both inhibitors had more modest effects on *B. pahangi*, and *T. muris* adult worms, their overall performance raises interest in better clarifying relative binding affinities to beta-tubulins from mammals and nematodes, effectiveness against benzimidazole resistant-parasitic nematodes, and potency among analogues that exist for each of these inhibitors [63–65].

### Advances on histopathologic methods and applications

Although the experimental focus here was on intestinal cells, DIC and bisbenzimide staining can rapidly provide information on most, or all, organs and tissues of the whole *A. suum* L3 and L4. While histopathologic sections provide obvious application to the research, we found *A. suum* L3 and L4 unexpectedly receptive to assessment by DIC microscopy of unfixed specimens and live staining by bisbenzimide (Hoescht 33258), a cell permeable DNA dye superior to the more commonly used DAPI for this purpose [66]. Although bisbenzimide stain may present a confounding factor during treatment, concordance between results from this assay and histopathology sections of non-bisbenzimide stained larvae greatly reduce this concern, and there was no indication of ill-effects in control larvae treated with bisbenzimide. Otherwise this live staining method has high value for monitoring many if not all nuclei among organ systems during larval development and in response to experimental treatments. The real time assessment capabilities supported by DIC and bisbenzimide provide important adjuncts to histopathologic analyses using fixed and sectioned material, and has potential application to numerous nematode species.

In conclusion, we have established a systems biology approach that integrates omics-based and chemogenomics-based predictive models to identify multiple inhibitors (prospective anthelmintics) with activity against phylogenetically diverse parasitic nematodes. These were coupled with methods that delineate pathologic profiles for each inhibitor that are based on multiple criteria (pathologic signatures) for application to multiple lines of future experiments. The approach reflects a first culminating step of a long-range design that integrates multiomics databases, evidence-based information and experimental methods focused on the nematode intestinal tract, to elucidate nematode intestinal toxicants with potential application to anthelmintic research. The general approach can be extended to additional cellular pathways identified in this research as well as multiple other tissues of parasitic nematodes.

## ACKNOWLEDGMENETS

We thank Susan Smart (WSU) for excellent technical support, and Dr. Allan Pessier (WSU) for insightful discussions. The research was supported by the National Institute of General Medical Sciences Grant R01GM097435 to M.M. The funders had no role in study design, data collection and analysis, decision to publish, or preparation of the manuscript.

## AUTHOR CONTRIBUTIONS

M.M and D.P.J. designed the study and directed the systems biology analysis. B.A.R. and R.T performed the data analysis. D.P.J., C.A.B. and J.S. performed the in vitro assays. B.B. and J. F. U. provided adult stage parasites and aided the in vitro assays. D.P.J., M.M., B.A.R and R.T. designed and prepared the illustrations and wrote the paper. All authors read and approved the final paper.

## MATERIALS AND METHODS

### Ethics Statement

All animal experiments were carried out under protocols approved by Washington State University Institutional Animal Care and Use Committee approved protocol 4097, the University of Missouri Animal Care and Use Committee approved protocol 9537) and United States Department of Agriculture the Institutional Animal Care and Use Committee (IACUC), approved protocol 18-029. Protocols meets requirements of AVMA Guidelines for the Euthanasia of Animals: 2013 Edition; Guide for the Care and Use of Laboratory Animals: 2011 Edition, National Research Council, and USA Animal Welfare Act and Animal Welfare Regulations: 2017 Edition (AWA), US Department of Agriculture.

### Prioritizing intestinal genes as potential anthelmintic targets

Since the most data was available for *Ascaris suum*, our intestinal gene target database was constructed for scoring targets based on the complete *A. suum* gene set [67]. Data from the other two core species (*Trichuris suis* and *Haemonchus contortus*) and from *C. elegans* was integrated in the dataset by identification of the best predicted protein sequence match to each predicted *A. suum* protein (using BLAST, *E*≤10^−5^). Prioritized candidates from this *A. suum*-based scoring system can be used to identify candidates across species using our comprehensive and high-quality orthologous group and intestinal expression database [23].

The prioritization (**Fig. 2**) consisted of target scoring based on four broad criteria, each of which had several individual scores assigned: A. Orthology - Only gene members belonging to Conserved Intestinal Families (cIntFams [23]; *A. suum*, *H. contortus* and *T. suis*) were considered (3,564 of the 18,542 total genes in *A. suum* gene set [67]). Genes are further prioritized based on orthology if they shared orthologs across all 10 nematode species (in order to prioritize conserved targets; score of 1 assigned if true; *T. suis, T. muris, T. spiralis, A. suum, B. malayi, L. loa, H. contortus, A. ceylanicum, C. elegans* and *N. americanus* [23]), and if they had low homology to host counterparts (in order to prioritize targets that have less of a possibility of disrupting host protein functions; score of 1 assigned if true); B. Intestinal proteomic evidence - being detected in a previously published *A. suum* intestinal proteomics study [21] (in any intestinal compartment, score of 1 assigned if true), and based on the level of detection in the intestine as quantified by spectral counts (scaled up to a maximum score of 1, using [number of spectra detected / 100]); C. Intestine expression level - Intestinal gene expression scores were calculated for *A. suum* (two different experiments [19, 28]), *H. contortus* and *T. suis* [23] according to 1-(expression rank after FPKM normalization / number of expressed genes). For *A. suum*, the average of the scores from the two datasets was used, and the scores for the other species (1 point each) were aligned to the *A. suum* genes according to the best sequence match to the protein sequences (BLAST, *E*≤10^−5^). The maximum total expression score between the three species was 3, prioritizing genes with high intestinal expression levels; D. Functional annotations - First, *A. suum* genes with a top-matched *C. elegans* ortholog that has a severe RNAi phenotype (e.g. lethal, sterile, intestine-specific phenotype, according to WormBase [30–33]) were assigned a score of 1, to prioritize targets with known desired phenotypes. Second, KEGGScan [68] (using KEGG release 78 [29]) was used to assign KEGG Orthologous Groups (KOs) and these were assigned to KEGG pathways for each protein; a score of 0 was assigned for proteins mapped to no pathways, and proteins in pathways were scored according to 0.5 + [0.1 * [Number of KEGG pathways the protein’s KO was annotated]] (maximum value of 1), where more pathways were scored higher in order to prioritize proteins with a high impact to cellular function if they are inhibited. Third, in order to reduce the possibility that proteins serve a redundant biological function (and would therefore not have a severe phenotype), proteins with unique KOs among the total protein set were scored higher according to 1 - [# of other genes sharing KO/10], with a minimum value of zero when 10 or more other proteins in the *A. suum* gene set share the function. Fourth, all protein-protein interaction data from the Worm Interactome Database [27] (version 8) was matched to *A. suum* based on the best *C. elegans* sequence match (as described above). Proteins with no predicted PPIs were assigned a score of 0, and other proteins received a higher score for being matched to more protein-protein interactions according to 0.5 + [0.05 * [Number of KEGG pathways the protein’s KO was annotated]] (maximum value of 1).

### Chemogenomic screening for small molecule inhibitor prioritization

Inhibitors were prioritized using available data from the ChEMBL [69] database (**Fig. 1**). ChEMBL [69] targets were annotated for all *A. suum* inferred protein sequences (blastp E-value ≤ 10^−10^). Hereafter, the term “inhibitor” will be used here to represent all prospective small molecule inhibitors based on our computational and experimental process, although not all of the compounds are FDA-approved drugs, and some may potentially act as activators rather than inhibitors. ChEMBL was then used to match inhibitors to the assigned targets, identifying target:inhibitor pairs with a pChEMBL score ≥ 5. The number of *A. suum* genes matched to each inhibitor was used to calculate the **scaled gene count score** (# / 50, maximum value of 1), for each inhibitor that had a “Quantitative Estimate of Druglikeness” (weighted QED) score [70], an **inhibitor property prioritization score** (scaled to a maximum value of 1) was calculated by adding (i) the weighted QED scores (scaled between 0 and 1) and (ii) a value of 1 if the inhibitor could be administered orally or topically.

### Pathway enrichment among prioritized intestinal genes

All of the 3,564 scored *A. suum* genes were ranked according to their gene prioritization scores, and this ranked list was used as input for Gene Set Enrichment Analysis (GSEA [71]) based on KEGG [72] pathways (annotated per gene using KEGGScan [68]; **Fig. 1B)**. This approach identified the KEGG non-metabolism and metabolism pathways (**S3 Table**) that were significantly enriched among higher-scoring gene targets. This approach allowed for the identification of the most biologically interesting inhibitor target pathways, independent of inhibitor information (which will be addressed in the following steps). Of particular interest was the most significantly enriched pathway, the exocytosis [73] / synaptic vesicle cycle pathway (ko04721, P<10^−10^), which contained 37 cIntFam [23] genes, many of which were high-scoring (**S4 Table**). This pathway had 5 members among the 50 top-ranking genes, which included 3 ATPases (GS_07654 [#3 ranked overall], GS_12676 and GS_02407), one clathrin heavy chain gene (GS_17518 [#6 ranked overall]), and one MFS transporter (GS_06670). Due to the large number of high-scoring genes, the coverage of strong inhibitor target candidates across the pathway, and the biological significance of this pathway in terms of parasite survival and host interactions [11, 74], these genes were prioritized for downstream drug targeting.

### Final prioritization of the top enriched pathway (exocytosis)

Based on the pathway enrichment analysis (**Fig. 1B**), only inhibitors matched to *A. suum* genes from the exocytosis KEGG pathway were included for the final prioritization. Additionally, inhibitors with an annotation of “NULL” in the ChEMBL database were also excluded since these are relatively untested and unstudied. After this filtering, the **final inhibitor prioritization score** was calculated (**Fig. 1D**) by multiplying the maximum matched **gene prioritization score** (to target the most biologically relevant *A. suum* genes), the **inhibitor prioritization score** (to choose inhibitors with properties likely to result in treatment success) and the **scaled gene count score** (to choose inhibitors which target multiple *A. suum* gene targets, for maximum potential effect). The top 25 inhibitors and their respective scores are shown in **S1 Table**.

To expand on targeting the exocytosis function, inhibitors targeting exocytosis-associated proteins were also selected for testing independent of the scoring approach. As one example, tubulin is the target of the highly efficacious benzimidazole anthelmintics [75], which disrupts microtubules, a critical component in exocytosis [76]. Among our highest-scoring tubulin genes was GS_01240 (score 8.7/11), that was (A) conserved across nematode species; (B) identified in the *A. suum* intestine by proteomics; (C) more highly expressed in the intestine than 97.3% of genes in all 3 model intestinal species; (D) predicted to have an “embryonic lethal” RNAi phenotype in *C. elegans* and (E) predicted to be druggable according to its best hit PDB entry. In previous research, its ortholog (ben-1) was identified as the only benzimidazole-sensitive beta-tubulin in *C. elegans* [77], highlighting the value of the prioritization system in *de-novo* identification of inhibitor targets. A second beta-tubulin gene (GS_23993; score 8.7) had similar properties to GS_01240. Although there are 37 predicted tubulins in the *A. suum* genome, our prioritization approach identifies those that are intestine-associated.

From the foregoing rational, two beta-tubulin inhibitors were included that either showed toxic effects against *C. elegans* (Taltobulin [78]) or is undergoing clinical trials for use in humans (Combretastatins [79]). To gain additional breadth in anticipated targets, a Rab GTPase inhibitor was included, CID 1067700 [80], because Rab GTPases are apparently involved in cycling of endosomal/exosomal vesicles in *C. elegans* intestinal cells [53, 54]. Staurosporine was also include and is a broad specificity kinase inhibitor, and inhibitor of protein kinase C/exocytosis [81, 82]. Staurosporine was previously found to have low IC_50_ levels of potency on parasitic nematodes, but without target tissues identified [38], and we wanted to determine if intestinal cells are one of the targets.

The thirteen inhibitors prioritized by both approaches (**Fig. 1**) are listed in **Fig. 2**, and their structures are provided in **S1 Fig.** The following are the suppliers used to obtain the inhibitors for performing the phenotypic screens: Alvocidib (S1230), Sunitinib (S7781), Selleckchem Houston, TX; CID 1067700 (SML054), Combretastatin A4 (C7744), Fasudil HCl (CDS021620), Leflunomide (L5025), Podophyllotoxin (Podofilox; P4405), Tofacitinib (PZ0017), Sigma-Aldrich, St. Louis, MO; KW2449 HCl (B1208), BioVision, Milpitas, CA; Ruxolitinib (tlrl-rux), InvivoGen, San Diego, CA; Staurosporine (S-9300), LC Laboratories, Woburn, MA; Taltobulin (HY15584), MCE Monmouth, NJ.

### *In vitro* inhibitors screening in lung stages of *A. suum*

All animal protocols were approved by the Washington State University Institutional Animal Care and Use Committee. To produce *A. suum* lung stage larvae, adult female *A. suum* were collected from the intestines of swine that were processed at the University of Idaho Meat Science Laboratory (Moscow, Idaho). Eggs were stripped from the last 3 cm of *A. suum* uterus, washed in PBS then decoated using 0.25% hypochlorite until decoating was observed to have occurred (usually within 4 minutes). Decoated eggs were rinsed in 50 mL double distilled water 3 times, and eggs were then cultured to the infective stage at 20°C for 60 days in 0.1 M H_2_SO_4_ [83]. Larvated eggs were then washed in 50 mL distilled water 3 times and stored at 4°C until used.

Third-stage larvae (L3) were obtained from lungs [84] and trachea of New Zealand white rabbits (5.5 to 6.5 weeks old, Western Oregon Rabbit Company, Philomath, OR) after oral infection with 2,000 to 4,000 larvated eggs. Intact lungs, including trachea, were dissected from euthanized rabbits at 8 days post-infection, and L3 obtained by first flushing the trachea with approximately five 1 mL aliquots of warm PBS (37°C, starting temperature), using a micropipettor (Gilson) and 1 mL micropipette tip. Larvae were aspirated into the pipette tip and pooled. Approximately 2.5 cm of trachea were then removed from the anterior end, allowing access to bronchi with the pipette tip, and PBS was lavaged into left and right lobes of the lungs. Larvae extracted with each volume were visible in the pipette tip. The entire process involved lavage of 1 mL aliquots for up to a total of about 25 mL. Lavage was ended when no more larvae were observed in lavage extract. L3 obtained in this manner were settled by gravity and then washed in 3 sequential 50 mL volumes of warm PBS followed by 3 sequential 15 mL volumes, with intervening gravity sedimentation and discard of supernatant PBS. The lavage method required about 1 hour to remove lungs and produce approximately 300 larvae from each rabbit, although the number of larvae varied somewhat among preparations. Extracted and cleaned larvae were then suspended in RPMI medium containing 10% swine serum, 100 units penicillin and 100 μg Streptomycin/mL (P0781, Sigma Aldrich, St. Louis MO) and then dispensed into wells of 96-well plates (Costar, Corning Inc., Corning, NY, triplicate wells for each treatment), with a total volume of 100 μL culture medium containing 1 μL of inhibitor treatment dissolved in DMSO, or 1 μL of DMSO alone in medium for control wells. L3 were then cultured at 37°C for 5 days in 5% CO_2_. When fewer than 5 larvae were dispensed in a well (6 times out of 369 wells (1.6%); all in L3 IC_50_ experiments), this was noted and results obtained were evaluated relative to adjacent time points and adjacent inhibitor concentrations. In no case were comparatively erratic outcomes observed.

L4 (fourth-stage larvae) were obtained by routine culture of L3s for 3 days (about 88% of L3 molted between days 2 and 3 in culture (see results) without treatments and with daily replacement of media. L4 were then dispensed into wells of 96-well plates for culture and experimental treatment under conditions identical to those used for L3. In other reports, lung stage L3 were collected on day 7 [46] and molting occurred 1 day later in culture (between days 3 and 4) than observed here. Thus, molting occurs in both systems around 10 to 11 days post-infection irrespective of when lung stage L3 are collected.

Motility of *A. suum* L3 and L4 was routinely assessed microscopically, but daily on days 1 through 5, using a Nikon Diaphot 300 inverted microscope equipped with a Nikon D5100 digital camera and epifluorescence capabilities. Otherwise immotile larvae that displayed an occasional twitch were considered immotile. In addition, treated *A. suum* L3 were scored for presence of shed cuticles, indicating molting to L4, but percentage shed was not quantified in all experiments. Other effects on morphology were noted, some of which were quantified as described in results. Effects of inhibitor treatments were expressed as mean percentage motile, or with a given morphology, compared to respective wells on day 0.

IC_50_ experiments were conducted on *A. suum* L3 and L4 using two-fold dilutions beginning with 500 μM to 31.25 μM final concentrations (5 treatment levels each in triplicate wells) for all inhibitors tested, except Staurosporine which ranged from 50 μM to 3.125 μM. Treatments were delivered in a 1 μL volume of DMSO.

### *In vitro* inhibitor screening in adult whipworm *Trichuris muris* and adult filarial worm *Brugia pahangi*

Adult *Trichuris muris* were removed from the cecum and proximal colon of infected C57Bl/6/STAT6 deficient mice (Beltsville IACUC protocol #18-029) using forceps between 32-35 days after inoculation with infective eggs and washed by sedimentation three times with media (RPMI-1640 with 25 mM HEPES, 2.0 g/L NaHCO_3_, 5% heat inactivated FBS, and 1X Antibiotic/Antimycotic solution). The works were incubated for 1-2hrs at 37C in a water bath and then washed again as above and shipped to UCSF overnight. On the day of arrival (day 0), adult worms were wash as described above plated into 24-well plates containing 500 µl media per well, with two worms per well. Based on IC_50_ values obtained in *A. suum* L3 and L4 were treated with 100 µM, except for Staurosporine, which was tested at 25 µM and 2.5 µM. Control worms were treated with 1% DMSO. Four replicate wells (8 worms total) were used per inhibitor and worms were maintained in a 37° C incubator with 5% CO_2_. Motility was measured using the Consensus Voting Luminance Difference algorithm WormAssay software as described by Marcellino et al. 2012 [85]. Motility readings were taken daily on days 0 to 6.

*Brugia pahangi* adult females were collected from male Mongolian gerbils (*Meriones unguiculatus*, Charles Rivers Labs) and incubated in media (RPMI-1640 with 25 mM HEPES, 2.0 g/L NaHCO_3_, 5% heat inactivated FBS, and 1X Antibiotic/Antimycotic solution) and maintained in a 37°C incubator with 5% CO_2_ overnight. The following day (Day 0), media was exchanged and worms were plated individually into 24-well plates containing 500 µl media per well. Worms were treated with 100 µM inhibitor, except for Staurosporine, which was tested at 25 µM. Control worms were treated with 1% DMSO and each inhibitor was tested with 4 replicates. Motility was measured using the Lucas-Kanade Optical Flow algorithm WormAssay software as described by Marcellino et al. 2012 [85]. Motility readings were taken daily on days 0 to 6. Results for both the *T. muris* and *B. pahangi* motility assays were reported as percent inhibitions based on the motility of their respective DMSO controls.

### *Brugia pahangi* L3 *in vitro* molt assay

*B. pahangi* L3 were collected from *Aedes aegypti* Liverpool (LVP) strain mosquitoes 13 days after infection via blood meal and shipped to UCSF overnight. On the day of arrival (day 0), L3 were washed 3X with wash media (RPMI-1640 + 1X Antibiotic/Antimycotic solution + 10 μg/mL gentamycin + 2 μg/mL ciprofloxacin), then washed once with culture media (MEM alpha with nucleosides [Gibco catalog #12571-063] + 10% heat-inactivated fetal bovine serum + 1X penicillin/streptomycin + 10 μg/mL gentamycin + 2 μg/mL ciprofloxacin + 2 μg/mL ceftazidime), plated into 96-well plates with about 5 larvae in 200 μL of culture media per well and were maintained in a 37°C, 5% CO_2_ incubator. On day 4 of culture, 100 μL of media was removed from each well and replaced with culture media containing the test inhibitor. The inhibitors were tested at the following concentrations: Sunitinib - 100, 62.5, 31.25, 15.6 and 7.8 μM; Staurosporine - 25 μM, Tofacitinib - 125 and 62.5 μM; Camptothecin - 62.5 and 31.25 μM and control worms were treated with 1% DMSO. On the following day, 100 μL of media was removed and replaced with culture media containing 30 μg/mL ascorbic acid (Sigma catalog #A4544) for a final concentration of 15 μg/mL [86] plus sufficient drug to maintain the concentrations listed previously. Motility was rated by visual examination on a scale of 0 (no movement) to 5 (fully active) on day 4 (before drug treatment), day 5 (before the addition of ascorbic acid), and days 7, 8, 9, and 12. Percent inhibition of motility was calculated by dividing the mean motility units of treated larvae by the mean motility of control larvae, subtracting this number from 1 and multiplying by 100%. Molting was measured by counting the number of casts present in the wells on days 8, 9, and 12. Percent inhibition of molting was determined by calculating the percentage of L3 that molted to L4 within each treated well, dividing this by the molting percentage of DMSO controls, and subtracting this number from 1 and multiplying by 100%. Prism (version 6.0f 2014, GraphPad Software, Inc) was used to calculate and graph the IC_50_s.

### *In vitro* inhibitors screening in *C. elegans*

The *C. elegans* N2 strain was used to test inhibitors for inhibition of movement and morphological effects. Synchronized larval populations initiated with eggs from adult worms were prepared by a standard protocol [87]. Eggs obtained by treatment of adult worms with 1% hypochlorite were washed 3 times in 4 mL of M9 medium and pipetted onto agar plates with lawns of OP50 bacteria and cultured overnight. Hatched larvae were collected by suspension in M9 media, pelleted and resuspended in M9 media, then dispensed into wells of 48 well plates (BioLite, Thermo Fisher Scientific, Rochester, NY) in a total volume of 100 μL, including 10 μL of OP 50 resuspended in M9 media from a pellet of a fresh 6-hour culture, and 1 μL of inhibitor solution in DMSO, or DMSO alone for control wells. A minimum of 5 larvae were included per well. Plates for 1- or 2-day old larvae received inhibitor treatments (in triplicate wells) and observations were made at 48 hrs post-initiation of cultures, as described for *A. suum* larvae. Motility was scored as movement or no movement (with agitation of the plate), along with notes of overall movement in a well compared to control worms. Otherwise immotile larvae that displayed an occasional twitch were considered immotile. Effects of inhibitor treatments were expressed as mean percentage motile compared to respective wells on day 0.

### Pathological effects in whole *A. suum* larvae, and intestinal cells and tissue

Endpoint morphologies of treated larvae were typically recorded after day 5 of treatment, but timing was adjusted as needed to capture relevant results, using a Nikon Diaphot 300 inverted microscope equipped with epifluorescence capabilities and a Nikon D5100 digital camera.

In addition, intestinal effects of selected inhibitors were evaluated on day 2 following treatment of L3 by 1) differential interference contrast (DIC) microscopy; 2) pre-treatment for two hours with the cell permeable nuclear stain Hoescht 33258 (bisbenzimide 10, µg/mL) prior to co-culture with inhibitors to assess effects on intestinal cell nuclei; or 3) hematoxylin and eosin stained histological sections (Histology laboratory of the Washington Animal Disease Diagnostic Laboratory, Pullman WA) of formaldehyde (3.7% solution) fixed L3 following inhibitor treatments. Visualization by DIC or of bisbenzimide stained worms was done on unfixed samples rinsed free of stains using phosphate buffered saline until background was negligible (usually 3 × 200 µL rinses). Results from each of the methods listed were obtained from independent experiments that, with exception of histopathology staining, were conducted at least three times.

Observation from DIC and bisbenzimide assessments were made using a Nikon Optiphot compound microscope equipped with DIC filters, epifluorescence capabilities and a Nikon D5100 digital camera. To optimize resolution, images were captured in movie mode, and then selected screen shots were copied and used to produce final digital images. Images of histological sections were recorded using an Olympus CX41 compound microscope with digital recording capabilities supported by DP manager and controller software. Standard blue, green and red fluorescence filters were used to capture fluorescence images.

### Statistics

For the *A. suum* and *C. elegans* assays, IC_50_s were estimated using R dose-response analysis[88]. For this analysis, the “mselect” function was used to select the best-fit dose-response model (using AIC criterion), with the lower and upper limits of efficacy constrained by specifying the corresponding parameters of the model (to 0 and 1, respectively). In all the cases, the best-fit model was Weibull function (W1.4 or W2.4). Significance of the fit was estimated using “neill.tes” by estimating P-value of lack-of-fit. **S3 Fig.** illustrates one of the cases with significant lack of fit. IC_50_s out of the range of screened concentrations only had the corresponding upper or lower bound reported. Significant pathway enrichment was tested using Gene Set Enrichment Analysis (GSEA [71]) based on KEGG [72] pathways (annotated per gene using KEGGScan [68]), and FDR correction to the P values was applied to correct for multiple testing. T-tests were performed using a two-tailed test with unequal variance, and one-way analysis of variance (ANOVA) testing was performed with a Tukey HSD post-hoc test.

## SUPPORTING INFORMATION

**S1 Fig.** Chemical structures of the 13 inhibitors studied.

**S2 Fig**. Overview of *A. suum* L3 molting assay methods and results.

**S3 Fig.** Motility response curve examples. (**A**) An example to illustrate a lack of fit of dose-response curve. Ruxolitinib Day 2 data (black) showed significant lack of fit for *A. suum* L3 larvae, primarily due to anomalously high motility of the 500 μM dosage samples. Data for Days 4 and 5 (red and green, respectively) showed good fit. (**B**) L3 Motility curves for Leflunomide (**1**). There is rapid inhibition of motility for 250 and 500 μM dosage, but concentrations below 125 μM show delayed inhibition, resembling effects of Sunitinib (**6**) and Tofacitinib (**10**) on L3.

**S4 Fig.**: *B. pahangi* molting phenotypes. (**A**) DMSO control L4 larvae, showing a successful molt. (**B**) Larvae with a bump/protrusion, observed in several of the larvae treated with 16 μM Sunitinib. (**C**) Example of an L3 that has failed to molt. This phenotype occurred in both treated and DMSO controls that fail to molt.

**S5 Fig.** Motility inhibition for 24 and 48 hours-old *C. elegans* larvae. Treatment responses for (**A**) all 13 inhibitors (1mM, except for Staurosporine at 100 µM), and motility was assessed after 48 hours of treatment. (**B**) 500 μM Leflunomide treatment and motility was assessed after 30 minutes of treatment. P values represent results from a two-tailed T-test (unequal variance).

**S1 Table**. The top 25 scored inhibitors. Tested inhibitors are indicated with an asterisk.

**S2 Table**. The top 50 genes ranked based on prioritization score

**S3 Table**. Top enriched metabolism and non-metabolism KEGG Pathways

**S4 Table**. All cIntFam genes belonging to the Exocytosis KEGG pathway (“synaptic vesicle cycle”, ko04721)

**S5 Table.** Selected characteristics of the thirteen experimentally tested inhibitors.

